# Relationship between Deceleration Morphology and Phase Rectified Signal Averaging-based Parameters during Labor

**DOI:** 10.1101/2021.04.21.440741

**Authors:** Massimo W. Rivolta, Moira Barbieri, Tamara Stampalija, Roberto Sassi, Martin G. Frasch

**Affiliations:** Dipartimento di Informatica, Università degli Studi di Milano, Milan, Italy; Unit of Fetal Medicine and Prenatal Diagnosis, Institute for maternal and child health IRCCS Burlo Garofolo, Trieste, Italy; Department of Medicine, Surgery and Health Sciences, University of Trieste, Trieste, Italy; Department of Obstetrics and Gynecology and Center on Human Development and Disability (CHDD), School of Medicine, University of Washington, Seattle, USA

**Keywords:** Phase-Rectified Signal Averaging, animal model, fetal heart rate, electronic fetal monitoring, HRV, fetal hypoxia, labor

## Abstract

During labor, uterine contractions trigger the response of the autonomic nervous system (ANS) of the fetus, producing sawtooth-like decelerations in the fetal heart rate (FHR) series. Under chronic hypoxia, ANS is known to regulate FHR differently with respect to healthy fetuses. In this study, we hypothesized that such different ANS regulation might also lead to a change in the FHR deceleration morphology. The hypothesis was tested in an animal model comprising 7 normoxic and 5 chronically hypoxic fetuses that underwent a protocol of umbilical cord occlusions (UCOs). Deceleration morphologies in the fetal inter-beat time interval (FRR) series were modeled using a trapezoid with four parameters, i.e., baseline *b*, deceleration depth *a*, UCO response time *τ*_*u*_ and recovery time *τ*_*r*_. Comparing normoxic and hypoxic sheep, we found a clear difference for *τ*_*u*_ (24.8 ± 9.4 vs 39.8 ± 9.7 s; *p <* 0.05), *a* (268.1 ± 109.5 vs 373.0 ± 46.0 ms; *p <* 0.1) and Δ*τ* = *τ*_*u*_ − *τ*_*r*_ (13.2 ± 6.9 vs 23.9 ± 7.5 s; *p <* 0.05). Therefore, the animal model supported the hypothesis that hypoxic fetuses have a longer response time *τ*_*u*_ and larger asymmetry Δ*τ* as a response to UCOs. Assessing these morphological parameters during labor is challenging due to non-stationarity, phase desynchronization and noise. For this reason, in the second part of the study, we quantified whether acceleration capacity (AC), deceleration capacity (DC), and deceleration reserve (DR), computed through Phase-Rectified Signal Averaging (PRSA, known to be robust to noise), were correlated with the morphological parameters. DR and DC correlated with Δ*τ* and *τ*_*u*_ for a wide range of the PRSA parameter *T* (max Pearson’s correlation *ρ* = 0.9, *p <* 0.05, and *ρ* = 0.6, *p <* 0.1, respectively). In conclusion, deceleration morphologies have been found to differ between normoxic and hypoxic sheep fetuses during UCOs. The same difference can be assessed through PRSA based parameters, further motivating future investigations on the translational potential of this methodology on human data.

## 1 Introduction

During labor, a fetus might suffer considerable stress due to uterine contractions, causing transient oxygen reduction and head compression, resulting in vagal and sympathetic stimulations. Nutrient deprivation, hypoxemia, hypoxia, acidemia and cardiovascular decompensation directly impact the autonomic nervous system (ANS) and thus affect the fetal heart rate variability (FHRV) [1, 2]. The cardiotocography (CTG) remains the best available proxy of ANS’ functional state through the analysis of fetal heart rate (FHR) and its variability. Considering that the standard processing of CTG series has been found poorly correlated to the relevant clinical outcomes, such as fetal brain injury or death, new FHR biomarkers are needed to better quantify the risk of fetal morbidity and mortality during labor [3, 4].

Phase-Rectified Signal Averaging (PRSA) analysis extracts quasi-periodic oscillations from HRV series and it is more resistant to non-stationarities, signal loss and artifacts [5] than conventional HRV analysis techniques, such as the well-known spectral analysis. It provides two measures that quantify the average cardiac acceleration (AC) and deceleration (DC) capacities from an inter-beat time interval series (RR). Practically, these measures quantify the average RR increase (or decrease) in milliseconds. When quantified on CTG signals or fetal RR series (FRR), AC and DC seem to perform better than the short term variation of FHR in identifying fetal growth restricted fetuses [6, 7, 8] and adverse outcome [9]. In a study of fetal sheep exposed to repetitive umbilical cord occlusions (UCOs), a model of uterine contractions during labor, we found that there was a high correlation between AC and DC and acid-base balance [10]; particularly, AC and DC progressively increased with the severity of the UCOs, suggesting an activation of ANS of healthy normoxic fetus exposed to acute hypoxemia.

In the same animal model, we recently observed that, at the beginning of each UCO, FRR adapted by a progressive increase (reduction in FHR) and quickly recovered when pressure was released. In order to quantify such adaptations, we modeled the FRR deceleration using a first-order exponential model, one of the possible models typically employed for system modeling tasks [11], for both response and recovery phases [10]. These models were characterized by time constants, describing the speed of FRR adaptation (the larger the time constant, the slower the adaptation) and we found that healthy normoxic fetuses had longer UCO response times than the time necessary to return to the baseline level [10], suggesting the presence of asymmetric trends in the series during labor.

Motivated by this observation, we also proved that dissimilarities in AC and DC values arise when asymmetric increasing/decreasing trends appear in the series [12], which seem to occur during labor. We thus introduced the deceleration reserve (DR), a new PRSA-based metric for the quantification of such asymmetry [12]. The DR is computed as the difference between DC and AC. Up to date, DR was tested on a near-term pregnant sheep model and human CTG recordings, obtaining promising results for distinguishing between normoxic and chronically hypoxic fetuses, and to detect fetal acidemia. Even though PRSA processes the FRR series in its whole entirety, it is reasonable to hypothesize that AC, DC and DR are deeply linked with the FRR adaptation time due to uterine contractions during labor.

In this study, we hypothesized that the different ANS regulation under chronic hypoxia might also lead to a change in the FHR deceleration morphology, as an result of the uterine contraction, and that the adaptation times would be different from those of healthy fetuses. The hypothesis was tested in an animal model comprising 7 normoxic and 5 chronically hypoxic fetuses that underwent a protocol of UCOs. Deceleration morphologies on the FRR series were modeled using a trapezoid with four parameters characterizing the adaptation times, baseline and deceleration depth. The parameters were compared between the two groups. We also quantified their correlation with biomarkers of acid/base balance. Assessing these morphological parameters during labor is challenging due to non-stationarity, phase desynchronization and noise. For this reason, in the second part of the study, we quantified whether AC, DC and DR were correlated with the morphological parameters. Given the fact that PRSA is more robust with respect to phase-desynchronization, a correlation might further support the opportunity of using AC, DC and DR in the clinical settings.

## 2 Materials and Methods

### 2.1 Animal model and FHR data

An established pregnant sheep model of labor was retrospectively analyzed. A comprehensive review on the pregnant sheep model and its translational significance for human physiology, in particular for studies of the ANS, can be found in [13]. The animal cohort comprised of nine normoxic and five spontaneously chronically hypoxic near-term pregnant sheep fetuses which underwent periodic UCOs mimicking uterine contractions during labor.

The animal and experimental models were described elsewhere [14]. Briefly, sheep fetuses were monitored over a 6 hours period during which a mechanical stimulation was applied to the umbilical cord by using an inflatable silicon rubber cuff. A baseline period of approximately 1 hour with no occlusion preceded the study.

After that, UCOs were delivered every 2.5 minutes and lasted for 1 minute. Three levels of occlusion strength, from partial to complete, were performed: mild (MILD, 60 minutes), moderate (MODERATE, 60 minutes) and complete (SEVERE, 60 minutes or until pH *<* 7.00 was reached). The stimulation protocol ended with a recovery period. During the stimulation protocol, pH, base deficit (BE) and lactate (hereafter referred to as “biomarkers”) were quantified by means of fetal arterial blood samples collected every 20 minutes.

Sheep fetuses were categorized as chronically hypoxic if O_2_Sat *<* 55%, as measured before the beginning of the UCO stimulation protocol. In this study, we refer to the two models as “normoxic” and “chronically hypoxic”, respectively. As per experimental protocol, both models showed a progressive worsening acidemia of the hypoxic status until pH *<* 7.00 was reached (see Figure 2 in [12]).

Fetal ECGs were collected by means of electrodes implanted into the left supra-scapular muscles, in the muscles of the right shoulder and in the cartilage of the sternum, and digitized at 1000 Hz. FRR series were automatically extracted from the fetal ECG [15].

In this study, we only considered the SEVERE phase of UCOs since FHR mostly changed during this condition.

### 2.2 FHR series preprocessing

A preprocessing similar to the one proposed in [12] was adopted for both datasets. Briefly, FRR series were analyzed to determine whether they were suitable for further analysis in terms of noise level, by excluding those recordings with more than 10% gaps during the SEVERE phases. Two normoxic fetuses were excluded from the analysis because of the high amount of missing beats. Furthermore, FRR intervals greater than 1500 ms (40 bpm) were labeled as artifacts and substituted with an equivalent number of beats (calculated dividing the length of each artifact by the median of the 20 nearby FRR samples). The reconstructed samples were used neither in the model fitting nor as anchor points in the PRSA analysis (in this latter case, however, they contributed to the selection of nearby anchor points).

### 2.3 Geometrical model of deceleration morphology and its fitting to FHR data

In this analysis, we quantified the average characteristics of the FHR response to UCOs in terms of baseline level, deceleration depth, time necessary to reach a steady condition of the FRR during both UCO stimulation and resting phase in the normoxic and hypoxic datasets. To do so, we used a simple model to describe the time evolution of FRR during each cycle of UCO and rest and extract the relevant information. The procedure took two steps. First, we time-aligned all the FRR segments of 150 s starting from the beginning of each UCO. Second, a piecewise linear model was fitted using a semi-automatic approach based on least squares. The model was as follows during UCO stimulation

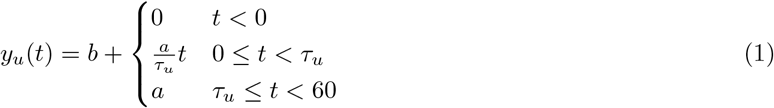

and the following one for the resting phase

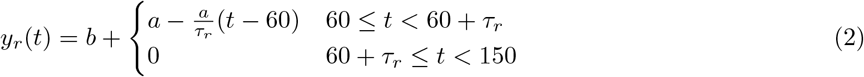

where *t* was the time (seconds), *b* (milliseconds), *a* (milliseconds), *τ*_*r*_ (seconds), and *τ*_*u*_ (seconds) were the morphological parameters to be estimated. In particular, *b* was the baseline FRR value in absence of UCOs, *τ*_*u*_ the time to reach the steady condition during UCO, *a* is the amplitude change of FRR, and *τ*_*r*_ the time to reach the baseline *b* after releasing the UCO. In addition, the difference Δ*τ* = *τ*_*u*_ − *τ*_*r*_ was considered as measure of asymmetry to the response to UCO stimulation. Figure 1 shows an example of model fitting and a visual description of the morphological parameters.

**Figure 1:**
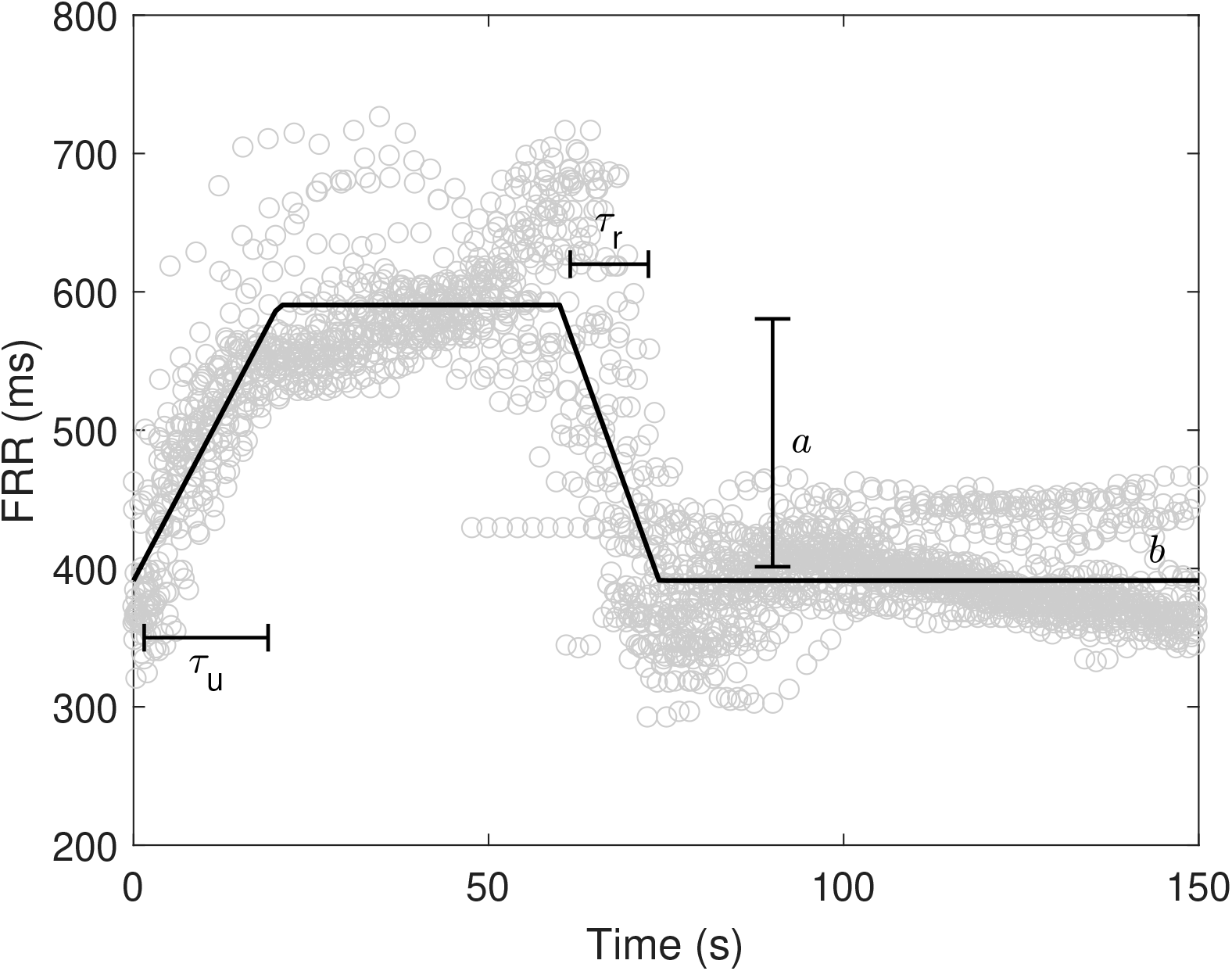
Example of model fitting along with variable definition. ° refers to FRR values from the beginning of the UCOs.

### 2.4 Correlation of deceleration morphology with time and biomarkers

In the second analysis, we determined whether the time intervals *τ*_*u*_, *τ*_*r*_ and Δ*τ* changed over time and were correlated with pH, base deficit and level of lactates, along the entire SEVERE phase. According to the stimulation protocol, blood samples were collected every 20 minutes up to the termination of the study. The same morphological parameters of eq. (1) and (2) were therefore estimated on all 20 minute windows preceding each blood sample. Two correlation analyses were thus performed. First, we computed the correlation coefficients between *τ*_*u*_, *τ*_*r*_ and Δ*τ*, and blood sample time points. Second, we determined the correlation between between *τ*_*u*_, *τ*_*r*_ and Δ*τ* with the biomarkers (pH, BE and lactate). To compensate for the fact that biomarkers’ values changed over time according to the stimulation protocol, partial correlation coefficients were computed, by accounting for the progress of time.

### 2.5 PRSA, AC, DC and DR

A complete description of the PRSA algorithm can be found in [5, 12]. The algorithm is divided into two steps. First, anchor points are identified on the time series *x*[*k*]. Each time index *k* that satisfies the condition

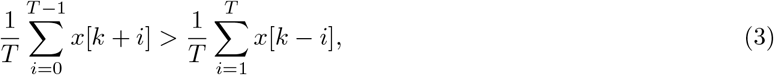

is inserted in the DC anchors’ point list (for AC, the inequality sign must be flipped). Second, all the windows of 2*L* elements centered on each anchor point are aligned (anchor points are located at the *L* + 1 sample) and then averaged. Such series of 2*L* averaged elements is the PRSA series.

From the PRSA series, AC and DC are then derived with

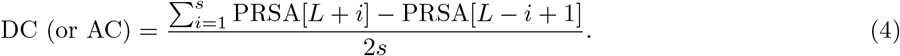

DR is instead defined as the sum of DC and AC (note that AC is a negative quantity for RR series) [12].

We quantified AC, DC and DR for multiple values of *T* (*s* = *T* and *L* = 50). A correlation analysis was performed to assess which range of *T* mostly correlated with the time constants derived from the piecewise linear models. In particular, we computed the correlation between *τ*_*u*_ and DC, *τ*_*r*_ and AC, and Δ*τ* and DR while varying the *T* value between 1 and 50.

Given the fact that: i) AC, DC and DR depend on the power of the series [12]; ii) a difference in the power of FRR signals was previously observed between the normoxic and hypoxic fetuses in this dataset [12]; and iii) a high variability in the deceleration depth *a* was observed (see sec. 3.1), we computed the partial correlation coefficients, while compensating for the amplitude *a* and baseline *b*. In this way, correlations were insensitive to linear relations of such quantities.

### 2.6 Statistical analysis

Results are reported as mean ± standard deviation and quantities were compared between the normoxic and hypoxic fetuses using a Student t-test. Correlations and partial correlations were computed using the Pearson’s correlation coefficient. Considering the low sample size, hypothesis tests and correlation coefficients were considered statistically significant when *p <* 0.1 (we also specify when *p <* 0.05).

## 3 Results

### 3.1 Comparison of morphological parameters between normoxic and hypoxic sheep fetuses

The morphological parameters obtained after model fitting were compared between the normoxic and hypoxic sheep fetuses. We obtained a model fitting achieving *R*^2^ values of 0.8 ± 0.1. Figure 2 reports the scatter plots for all pairs of morphological parameters for both animal models. A clear difference was found for *τ*_*u*_ (normoxic vs hypoxic; 24.8 ± 9.4 vs 39.8 ± 9.7; *p <* 0.05), no difference for *τ*_*r*_ (11.6 ± 4.8 vs 16.0 ± 3.9; *p >* 0.1), a difference in FRR change *a* (268.1 ± 109.5 vs 373.0 ± 46.0; *p <* 0.1), and no difference for the baseline *b* (357.0 ± 34.1 vs 372.6 ± 23.6; *p >* 0.1). Δ*τ* was found different between normoxic and hypoxic fetuses (13.2 ± 6.9 vs 23.9 ± 7.5; *p <* 0.05).

**Figure 2:**
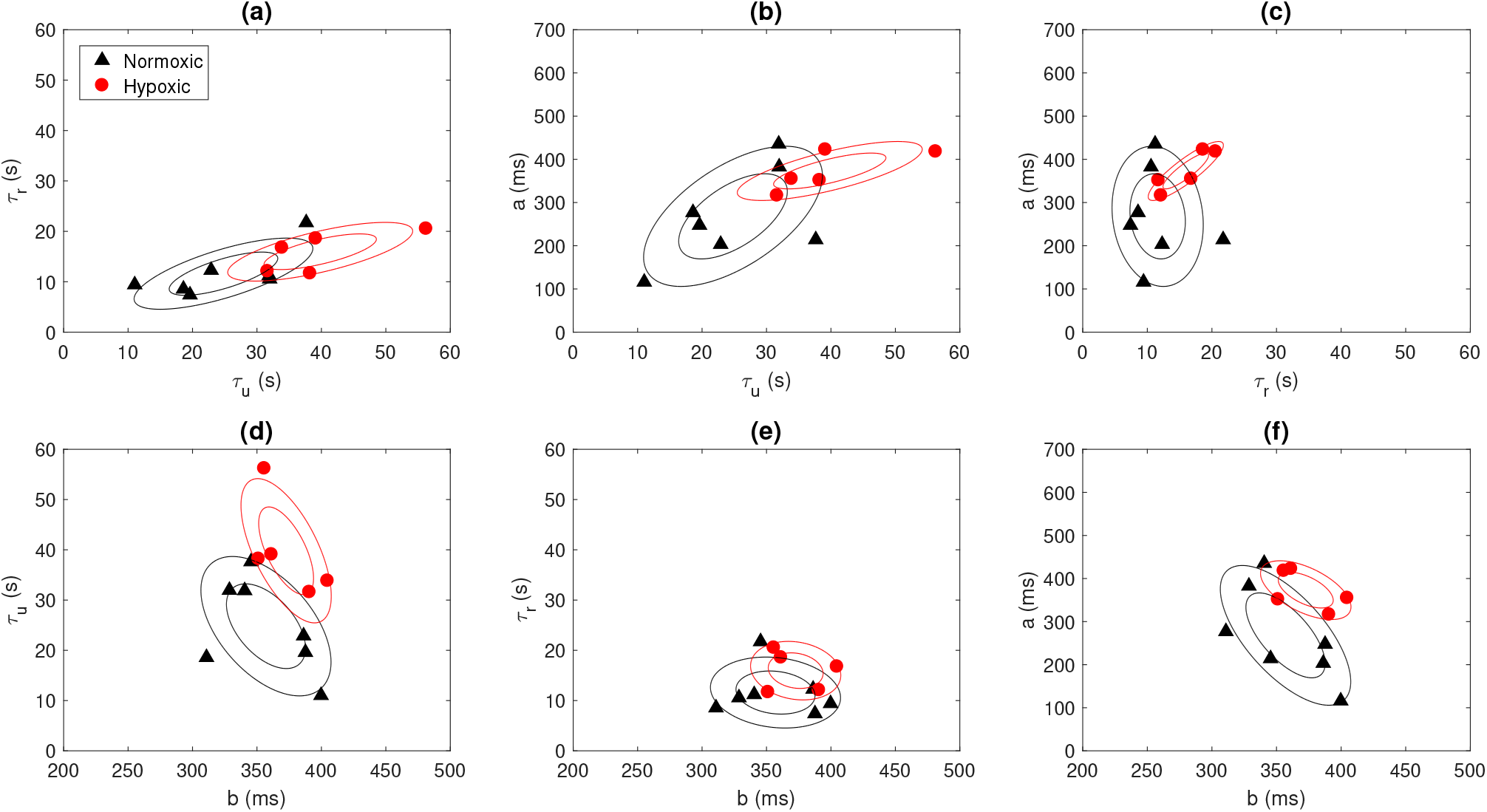
Scatter plot of the morphological parameters, *i*.*e*., *τ*_*r*_, *τ*_*u*_, *b* and *a*, for both normoxic and hypoxic fetuses. Contour plots are also reported (prepared with an assumption of Gaussianity, made for visualization purposes).

### 3.2 Time progression of FHR deceleration morphology and correlation with biomarkers

A correlation analysis was performed to assess the time progression of the morphological parameters over the entire duration of the SEVERE phase. We found no correlation between the morphological parameters and time for both animal models when considered together (correlations between time vs *τ*_*u*_, *τ*_*r*_, and Δ*τ* were −0.2, −0.1, and −0.3, respectively; *p >* 0.1) or separated (normoxic fetuses: −0.1, 0.1, and −0.1 and hypoxic fetuses: 0.1, 0.3, and −0.1; *p >* 0.1). Non-significant correlations were likely due to the limited sample size. In fact, a similar analysis performed on the biomarkers resulted in a moderate (significant) correlation with time (pH vs time: −0.5, *p <* 0.05; BE vs time: −0.6, *p <* 0.05; lactate vs time: 0.34, *p <* 0.1).

Consequently, when correlation was assessed between morphological parameters and biomarkers, partial correlation coefficients were calculated to account for this possible time variation. Partial correlations were found statistically significant for pH vs *τ*_*u*_ (−0.5; *p <* 0.05) and pH vs Δ*τ* (−0.6; *p <* 0.05), and for base deficit vs *τ*_*u*_ (−0.6; *p <* 0.05) and base deficit vs Δ*τ* (− 0.7; *p <* 0.05), whereas no significant correlation was found for lactate. Such significant correlations were mostly due to the normoxic dataset. Indeed, pH and base deficit were found correlated with the morphological parameters only for the normoxic data (*p <* 0.05). On the other hand, lactate was found correlated with only the morphological parameters of the hypoxic fetuses (coefficients for *τ*_*u*_ and Δ*τ* were −0.8 and −0.8; *p <* 0.05).

### 3.3 Correlation analysis for the PRSA parameters

Partial correlation coefficients were calculated between the PRSA and morphological parameters. A wide range of statistically significant correlations was found for DC vs *τ*_*u*_ (28 ≤ *T* ≤ 50; *p <* 0.1), reaching approximately −0.6 for large values of *T*. AC was not found correlated with *τ*_*r*_ for any value of *T* (*p >* 0.1). DR was found correlated with Δ*τ* for a wide range of *T* (4 ≤ *T* ≤ 50) with a maximal correlation at *T* = 9 of −0.9 (*p <* 0.05). An example of FRR after compensating for differences in *a* and *b* parameters, for both animal models, is shown in Fig. 3. A slower FHR adaptation, as response to UCO, becomes clearly visible for the hypoxic fetus after such compensation.

**Figure 3:**
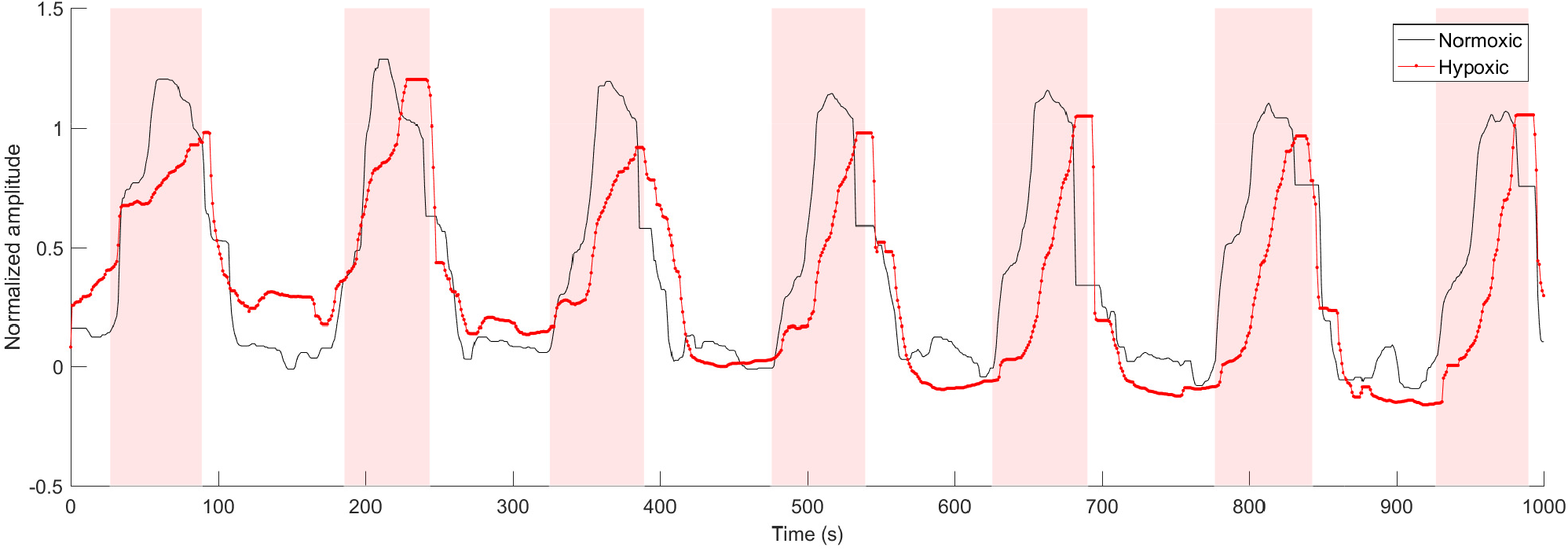
Example of normalized FRR signals from one normoxic and one hypoxic fetus (a median filter was applied to the signals to enhance the trend). The shaded area corresponds to UCOs.

**Figure 4:**
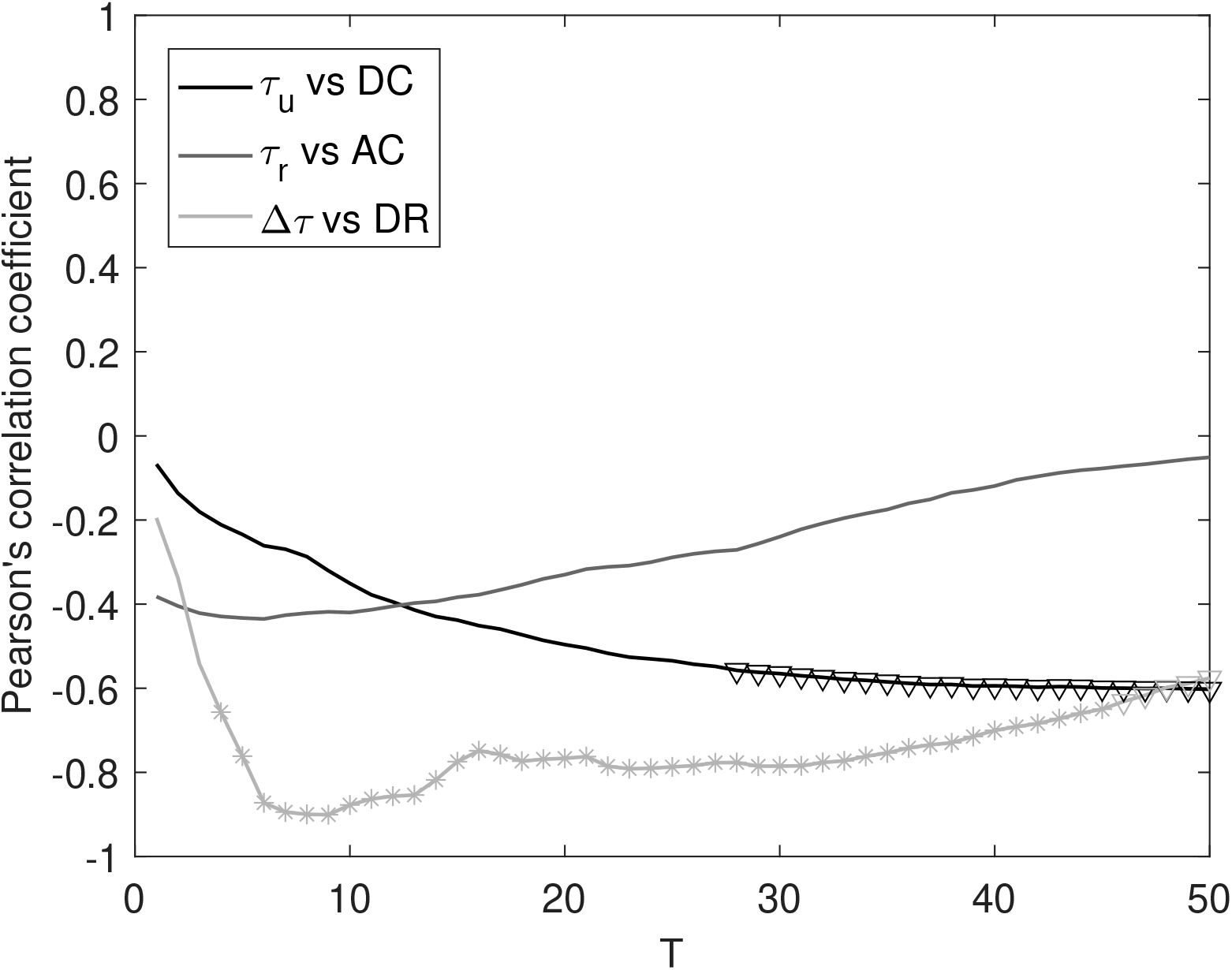
Partial correlation coefficient between *τ*_*u*_ vs DC, *τ*_*r*_ vs AC and Δ*τ* vs DR, for different values of *T*. * (*p <* 0.05) and ▽ (*p <* 0.1) refer to statistically significant correlations.

## 4 Discussion

### 4.1 Morphological differences in FHR decelerations

When comparing the morphological differences of FHR decelerations, the significant differences in *τ*_*u*_, Δ*τ* and *a* between normoxic and chronically hypoxic fetuses suggest a different FHR response to the ANS stimulation caused by UCOs in hypoxic fetus. These findings are in line with other reports of different response to external hypoxic stimulation, as in our case by UCOs, in normoxic versus already hypoxic fetuses [16]. The reduction of FHR in the presence of hypoxic stimuli in a normoxic fetus represents a protective mechanism while it reduces the oxygen consumption via reduced myocardial work [17]. In the presence of already established acidemia, the ANS modulation changes [18, 19], and some of the adaptive mechanisms, such as chemoreceptor mediated circulatory adaptation, might be altered due to progressive tissue damage, including brain damage. In this view, the fact that the correlation between biomarkers (pH and BE) and time was driven mainly by the normoxic fetuses should not be surprising. Both pH and BE are strong stimulators of chemoreceptors in normoxic fetus [20] and in our previous work we found a stronger correlation between AC and DC and pH and BE in normoxic fetuses exposed to UCOs, than with lactate concentration [10]. Thus, we speculate that the absence of the correlation between pH and BE in hypoxic fetus might represent a sign of already altered ANS and chemoreceptor function. Certainly, it has to be acknowledged that the number of available measurements was different for normoxic and hypoxic fetuses (21 vs 9), resulting in a much shorter SEVERE phase for the hypoxic ones. On the other hand, we found that the lactate correlated only with the morphological parameters in the hypoxic fetuses. A possible explanation could be due to the fact that the lactate represents the end product of anaerobic glucose metabolism, reflecting, thus, the metabolic acidosis [21]. Indeed, it has been suggested that lactate concentrations, on fetal scalp and umbilical artery at birth, might be a better predictor of poor neonatal outcome than pH [22].

### 4.2 Correlation between morphological parameters and AC, DC and DR

Morphological parameters Δ*τ* and *τ*_*u*_ were found to correlate with DR (4 ≤ *T* ≤ 50, with maximum at *T* = 9) and DC (*T* ≥ 28), respectively. In our previous study on the same animal model, we found that DR achieved the highest discriminatory power in distinguishing between normoxic and hypoxic sheep fetuses during the SEVERE phase of the protocol, with *T* ranging between 5 and 9 [12]. Such high discriminatory power was likely related to the larger Δ*τ* measured on hypoxic fetuses in this study. In other words, we found that the ANS regulation of already hypoxic fetuses during labor affects the deceleration morphology (particularly Δ*τ*), thus further highlighting that the presence of asymmetric trends in the series is relevant for risk stratification.

Correlations were not statistically significant for the entire range of *T* values considered. This was an expected result because the *T* value acts as frequency selector, specifically as a band-pass filter [23], whose frequency band shrinks when *T* increases. Although there is no clear evidence about the optimal *T* value for the detection of already hypoxic fetuses, previous studies employed effectively, for the detection of intra-uterine growth restriction (IUGR) during antepartum fetal monitoring, values of *T* corresponding to the range 2.5 s to 10 s [24, 6, 9]. On the other hand, fetal acidemia occurring during labor seems better detected at lower time scales between 0.5 s to 1.25 s [25, 12], thus suggesting a different mechanism for healthy fetuses during acute stress.

The PRSA series is also amplitude-dependent. In our previous study, we found a perfect linear relationship between the standard deviation of the series and the PRSA parameters [12] for Gaussian processes during stationary condition (*e*.*g*., a situation likely occurring during antepartum fetal monitoring). A similar relation is expected for other indices of variability, such as the short-term variation (STV). In fact, considering results obtained during fetal monitoring of IUGR fetuses, Huhn *et al*. [24] and Graatsma *et al*. [26] found a correlation of about 0.7 between STV and AC for IUGR fetuses antepartum. On the other hand, given the non-stationary nature of FHR series during labor and the fact that the PRSA algorithm is applied to the entire recording, the relationship between STV and PRSA may break, as supported by the study of Georgieva *et al*. [25] who reported a significant correlation of about 0.3 during labor. It sounds therefore reasonable that the long-term variability of FHR series may better correlate with PRSA parameters during labor. In fact, the deceleration depth *a* affects the variability of the entire FHR series (the standard deviation of the FHR series is proportional to *a*), and in turn, it affects the values of AC, DC and DR. However, while *a* was found larger in the hypoxic dataset, it is still unclear whether such morphological parameter would turn out to be important for risk stratification or needs to be considered as a confounding factor.

### 4.3 Comparison with Deceleration Area

The results reported so far are in line with other attempts of “capturing” the deceleration morphology for risk stratification. For example, the well-known “Deceleration Area” (DA) quantifies the severity of the deceleration taking into account both its depth and duration (the number of “missed” beats due to the deceleration). In the work of Cahill *et al*. [27], DA was quantified by approximating the deceleration by a triangle having a base as long as its duration and height corresponding to the depth, and then computing its area. DA was found to perform well for detecting fetal acidemia (AUC = 0.76). Using the morphological parameters, which are depicted in Fig. 1, DA is then given by

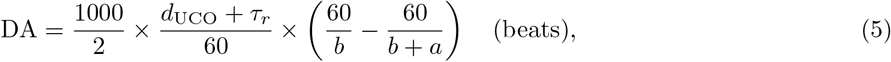

where *d*_UCO_ = 60 s is the duration of the UCO, and 1000 is the conversion factor from milliseconds to seconds. In our case, given the severity of the stimulation protocol, UCOs caused FHR responses which were better approximated by a trapezoidal model than a triangle. Thus, it was possible to calculate DA using also the following formula

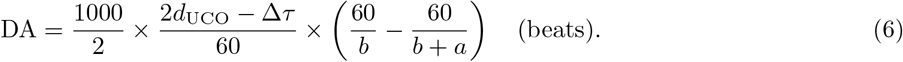

From these formulas, the relationship between DA and the morphological parameters, which we studied in this paper, is evident. The second formula also points out a link between DA and the asymmetry value Δ*τ* = *τ*_*u*_ − *τ*_*r*_, which is not captured in the triangular approximation usually employed. The relationship with Δ*τ* hints the importance of looking at asymmetric trends in the series present during labor. We leave this point to future investigations.

In our study, however, DA was not found significantly different between normoxic and hypoxic fetuses in our data (62.9 ± 19.5 vs 64.4 ± 6.6 beats; *p >* 0.1). A possible explanation could be the fact that Δ*τ* and *a* were higher for the hypoxic fetuses, and such quantities correlated with DA in opposite directions, thus making DA values indistinguishable due to balancing effects and (likely) the limited sample size. Another possible reason is that chronic hypoxia (in this study) and acidemia at birth [27] might trigger different regulatory ANS responses.

### 4.4 Limitations of the study

First, the sample size is small, dictated by the complexity of the animal model. Second, UCOs implemented in the experiments do not necessarily generalize to human labor, where the contractions are not equally regular nor are they all producing a complete occlusion of the umbilical cord. Third, sheep were analyzed during complete UCOs. However, changes in ANS activity in response to UCOs also occur earlier in time, when the UCOs are less severe or the recovery time between the UCOs is longer, and may reflect differences in the chronically hypoxic fetuses compared to the normoxic ones. Identifying these potentially earlier differences will be the subject of future studies. Fourth, all sheep fetuses displayed an individual pattern of pathological hypotensive responses to UCOs with regard to the timing of its emergence, with hypotensive responses to FHR decelerations showing well ahead of the severe UCOs in some instances [28, 29]. As our present study focused on the differences between the hypoxic and normoxic fetuses in the severe stage of UCOs, it did not investigate the relationship between the PRSA-based metrics and the timing of the onset of pathological hypotension. We leave this to future work.

### 4.5 Conclusions

Our study motivates further investigations on PRSA-related quantities to determine their potential advantage for risk stratification. It might also open interesting scenarios for interpreting PRSA-based results and improving FHR monitoring. The evaluation of the performance of these new metrics in identifying compromised fetuses during labor is still underway.

## Conflict of Interest Statement

MGF has a patent pending on abdominal ECG signal separation for FHR monitoring (WO2018160890).

## Author Contributions

MWR designed and implemented the analyses, and he drafted the manuscript. MGF collected the data. MWR and RS optimized the proposed mathematical framework. MB, TS and MGF were involved in the clinical interpretation of the results. All authors read, revised and approved the final manuscript.

